# An optimized protocol for *Candida albicans* infection in *Schmidtea mediterranea* to study fungal pathogenesis and host defense

**DOI:** 10.64898/2025.12.06.691972

**Authors:** Nora M. Shamoon, Beryl N. Arinda, Simriya Sandhu, Deepika Gunasekaran, Néstor J. Oviedo, Clarissa J. Nobile

## Abstract

*Candida albicans* is a common opportunistic fungal pathogen that asymptomatically colonizes most humans. Although typically a benign commensal, dysbiosis caused by antibiotic use, immune dysfunction, or epithelial barrier disruption can trigger fungal overgrowth and infection, ranging from superficial mucosal disease to life-threatening systemic candidiasis. New preclinical infection models are needed to dissect *C. albicans* pathogenesis *in vivo* across distinct stages of infection and with different measurable host outcomes. We previously established the planarian *Schmidtea mediterranea* as an invertebrate host for studying host-pathogen interactions during *C. albicans* infection. *S. mediterranea* relies entirely on conserved innate immune mechanisms capable of overcoming infection with pathogenic microorganisms, including bacteria and fungi. Planarians’ remarkable regenerative capacity and accessible stem cell populations make this organism a tractable model to analyze early immune responses, tissue repair, and pathogen clearance *in vivo*. This model supports simultaneous analysis of fungal virulence and host transcriptional responses, providing valuable insights into infection dynamics. Here, we present an updated protocol with detailed modifications, standardized procedures, and optimized steps for infecting *S. mediterranea* with *C. albicans*, designed to enhance reproducibility and enable systematic studies of fungal pathogenesis and host defense.

**SUMMARY:** We present an optimized and detailed protocol for using the planarian *Schmidtea mediterranea* as a model system to study host-pathogen interactions during fungal infection. This method builds on our previous procedure for infecting planarians with the human fungal pathogen *Candida albicans*, providing detailed guidance to enhance reproducibility and experimental consistency. The system enables comprehensive, parallel analysis of both host and pathogen responses throughout infection – from initial colonization and disease progression to diverse host outcomes, including clearance or mortality.

## INTRODUCTION

*Candida albicans* is a common opportunistic fungal pathogen that asymptomatically colonizes 50-70% of humans^1–3^. Typically, a harmless commensal, *C. albicans* can overgrow under conditions such as microbiota imbalance, antibiotic use, immunosuppression, epithelial barrier disruption, or the presence of medical devices, leading to diseases that range from superficial mucosal infections to life-threatening systemic candidiasis^4,5^. In immunocompromised individuals – including those with HIV/AIDS, patients undergoing chemotherapy or immunosuppressive therapies, individuals with chronic illnesses, or those in intensive care units – mortality rates can exceed 70%, underscoring the significant public health burden of invasive candidiasis^6–8^. Globally, invasive fungal infections affect more than 6.5 million people annually, with direct healthcare costs in the United States alone exceeding $8 billion per year for *Candida*-related infections ^9,10^. Despite available antifungal treatments, morbidity and mortality rates associated with *C. albicans* infections have remained largely unchanged for decades, highlighting the urgent need to better understand host-pathogen interactions in fungal disease.

*C. albicans* employs a diverse repertoire of virulence factors that support colonization, persistence, and pathogenesis across a range of host environments. These include adhesion to host tissues, immune evasion, cell-wall remodeling, secretion of hydrolytic enzymes, morphological plasticity, and biofilm formation^12–17^. Collectively, these traits allow *C. albicans* to adapt to diverse host niches, modulate immune responses, and resist antifungal therapies^17–22^. Elucidating how these processes shape infection outcomes – including host morbidity, mortality, and pathogen clearance – is essential for developing more effective preventive and therapeutic strategies.

To investigate these host-pathogen dynamics, we established a model system using the planarian *Schmidtea mediterranea* as a host^23–25^. We previously introduced this model as a versatile platform for studying the multisystem host response to fungal infection, leveraging a soaking-based infection method that enables synchronous exposure of animals to *C. albicans*. Here, we present updated step-by-step procedures that refine this protocol, improve reproducibility, and facilitate consistent implementation across laboratories.

Planarians are free-living invertebrates with exceptional regenerative capacity, capable of rapidly replacing tissues lost to injury or infection^26^. They lack an adaptive immune system and rely entirely on conserved innate defenses – including pattern recognition receptors, antimicrobial peptides, mucus secretion, and phagocytic cells – to clear bacterial and fungal pathogens within days^27–30^. Because innate immunity represents the first line of defense across eukaryotes and remains understudied outside mammalian systems^28^, planarians offer an opportunity to examine these processes *in vivo* with cellular and molecular resolution. Their regenerative ability is driven by adult pluripotent stem cells, known as neoblasts, which constitute approximately 30% of adult cells and contribute to recovery after infection^24,31^. These features make *S. mediterranea* a powerful model for dissecting pathogen virulence and host defense at genetic, cellular, tissue, and organismal scales, enabling us to address questions that are often difficult to study using traditional mammalian systems.

In addition, planarians’ small size (typically a few millimeters in length), low cost, and ease of maintenance^32^ support large-scale experiments while avoiding the financial, ethical, and regulatory constraints inherent to vertebrate models. Like *C. albicans*, *S. mediterranea* is genomically tractable, with a fully sequenced and annotated genome, and it supports advanced molecular and cellular techniques – including transcriptional profiling, highresolution immunohistochemistry and histology, and robust RNA interference (RNAi)^33–39^. These tools enable parallel analysis of host and pathogen responses during infection.

Several alternative preclinical models have been used to study fungal infections. Invertebrate hosts such as *Galleria mellonella* (wax moth), *Caenorhabditis elegans* (roundworm), and *Drosophila melanogaster* (fruit fly), a well as the vertebrate *Danio rerio* (zebrafish), each provide distinct experimental advantages but also have notable limitations. These systems often rely on survival as the primary endpoint because infections rapidly induce host death, preventing detailed assessment of morbidity, recovery, and multisystem responses – features that the planarian model readily captures.

Mammalian models such as *Mus musculus* (mouse), *Rattus norvegicus* (rat), *Oryctolagus cuniculus* (rabbit), and *Cavia porcellus* (guinea pig) more closely recapitulate human physiology but are limited by high costs, small cohort sizes, ethical and regulatory requirements, and the challenge of distinguishing innate from adaptive immune responses. Moreover, decades of fungal pathogenesis research in mice have focused predominantly on late-stage systemic infection^40–44^, often overlooking early events such as epithelial barrier disruption and mucosal overgrowth – the most common routes of fungal disease initiation in humans^45–47^.

Studying host-pathogen interactions is therefore essential for understanding the biological processes that underlie fungal virulence, host defense, and disease progression. Invasive fungal infections represent an escalating global threat, with multiple species now exhibiting resistance to all major antifungal drug classes. More than one billion people are estimated to be affected by fungal infections worldwide, and factors such as climate change, widespread antimicrobial use, and increasing numbers of immunocompromised individuals are accelerating the emergence of antifungal-resistant pathogens^48–51^. Understanding how pathogenic fungi colonize and engage their hosts is critical for developing new preventative and therapeutic strategies.

Here, we present a revised and standardized systemic infection protocol for *S. mediterraneaC. albicans* interactions, designed to promote reproducibility and to enable systematic investigation of fungal pathogenesis and host defense mechanisms. **Table 1** summarizes the key updates incorporated into this revised protocol.

**Table 1.** Key revisions to the *C. albicans*–*S. mediterranea* infection protocol.

|  |  |
| --- | --- |
| **System Feature** | **Updated Guidance** |
| Microbial culture | Prior to inoculation, remove all growth medium and wash *C. albicans* cells thoroughly with planarian water to eliminate residual medium. |
| Well volume | Reduce total well volume to 2,000 μL to increase the likelihood of planarian-*C. albicans* interactions. |
| Re-inoculation | Gently resuspend settled *C. albicans* cells in each well every 24 h using a sterile pipette to maintain consistent exposure throughout the infection period. |
| Specified animal inclusion criteria | Use size-standardized planarians (~5 mm in length). Exclude individuals with pre-existing damage, such as abnormal pigmentation, lesions, or developmental defects to ensure experimental uniformity. |
| ID<sub>50</sub> determination | Begin recording host outcomes at 1 day post-infection (dpi). Confirm deaths no earlier than 4 dpi to avoid misclassification of phenotypes, such as animals that appear moribund at 3 dpi but recover at later timepoints. |
| LD<sub>50</sub> determination | Record host survival and symptomatic data beginning at 1 dpi, as mortality can occur rapidly and not at a fixed timepoint. |
| Effective dose determination | Because effective infectious doses vary between planarian colonies, empirically determine the ID<sub>50</sub> and LD<sub>50</sub> for each planarian colony using dose response curves. Perform at least three independent replicates for statistical robustness. |
| Planarian colony maintenance | Planarian susceptibility is influenced by environmental conditions and maintenance practices. Feed and clean colonies regularly to maintain health and consistency. Reassess effective infectious doses following major personnel or husbandry changes. |

### PROTOCOL

This protocol builds on our previous methods and establishes clear, standardized procedures to enhance reproducibility and enable systematic investigation of fungal pathogenesis and host responses in *S. mediterranea*. The key refinements include prespecified animal inclusion and exclusion criteria, standardized microbial preparation, and comprehensive assessment of host endpoints. All refinements are summarized in **Table 1**.

The workflow provides detailed guidance for culturing *C. albicans*, infecting planarians through soaking-based exposure, and assessing fungal virulence and host responses using both qualitative and quantitative readouts across symptomatic and lethal-dose conditions. The optimized procedures presented here explicitly incorporate multiple variables that we have empirically identified as critical determinants of infection dynamics and host outcomes.

Step-by-step experimental procedures, data-analysis workflows, and scoring rubrics are provided below. All reagents, critical equipment, and media recipes are listed in the **Table of Materials** and **Supplementary File 1**.

**NOTE:** All procedures must be performed using appropriate sterile and aseptic biosafety level 2 techniques. Standard laboratory PPE – including gloves, lab coat, closed-toe shoes – should be worn at all times. Wash hands thoroughly after all experiments.

#### 1. Fungal Culturing

***C. albicans* Strain Information**

- *C. albicans* strain SN250 was used as the wildtype strain for all experiments.
- SN250 is derived from the clinical isolate SC5314, originally collected from a blood culture of a patient with disseminated candidiasis^52^.
- *C. albicans* strains are maintained as 25% glycerol stocks at −80°C.

##### Culturing *C. albicans* Strains for Infection

**Streaking from Cryogenic Stock**

1. Using a sterile applicator, streak the desired strain from −80°C stock onto YPD (1% yeast extract, 2% peptone, 2% glucose; pH 6.8) agar plates.
2. Incubate statically at 30°C for 72 hours.

**Preparing Overnight Cultures**

3. Inoculate a single colony into 4 mL liquid YPD in sterile borosilicate tubes (18×150 mm) using a sterile applicator.
4. Include a negative control tube containing uninoculated YPD under identical conditions.
5. Loosely cap the tubes to allow aeration and incubate overnight (16 h) at 30°C with shaking (∼220 rpm).

NOTE: If microbial growth occurs in the negative control, abort the experiment and restart with new media and sterile technique.

**Harvesting and Washing Cells**

6. Using aseptic technique, transfer cultures to labeled 15 mL centrifuge tubes.
7. Centrifuge cultures for 5 min at ∼3,000 × g to pellet the cells.
8. Carefully remove the YPD supernatant without disturbing the pellet.
9. Resuspend the pellet in planarian water (1× Montjuïc Salts: 1.6 mM NaCl, 1.0 mM CaCl₂, 1.0 mM MgSO₄, 0.1 mM MgCl₂, 0.1 mM KCl, 1.2 mM NaHCO₃ in Milli-Q water, pH 7.5) in a volume equal to that of the YPD used. *Example:* If combining 15 mL total of overnight culture, resuspend in 15 mL planarian water.
10. Mix thoroughly by vortexing.
11. Repeat steps 6-9 twice more to remove the growth media entirely.
12. After the final wash, resuspend the cell pellet in fresh planarian water.

##### Quantifying *C. albicans* Cell Concentration

**Dilution for OD Measurement**

1. Prepare three sterile 1.7 mL Eppendorf tubes with a 1:20 dilution (50 µL culture + 950 µL planarian water) or an alternative chosen dilution factor. Mix thoroughly.
2. Prepare a blank tube containing 1000 µL planarian water. **Spectrophotometry**
3. Set the spectrophotometer to measure optical density at 600 nm (OD₆₀₀).
4. Blank the spectrophotometer with planarian water.
5. Measure OD₆₀₀ for each diluted sample and record values.
6. Average the three readings and multiply by the dilution factor to determine the OD₆₀₀ of the stock culture.
7. Convert OD₆₀₀ to cell concentration NOTE: The optical density conversion factor should be determined empirically for each spectrophotometer. Based on our spectrophotometer, an OD₆₀₀ of 1 ≈ 2×10⁷ *C. albicans* cells/mL^53^.

**Determining Experimental Dosages**

8. Prepare the desired inoculum concentration for each experimental condition. NOTE: Perform at least three biological replicates to determine the effective dose for the planarian colony, particularly for symptomatic or lethal endpoints. Example Calculation (Symptomatic Endpoint)
• Observed ID₅₀ ≈ 7.0×10⁷ cells/mL for wildtype SN250
• Observed LD₅₀ ≈ 8.0×10⁷ cells/mL for wildtype SN250 Given a stock culture of 1.8×10⁸ cells/mL, calculate the dilution for a target of 8.0×10⁷ cells/mL in a final volume of 2 mL:

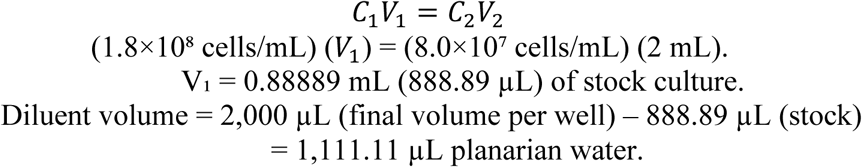

#### 2. Planarian Colony Preparation

##### Planarian Species and Maintenance

The CIW4 asexual clonal line of *S. mediterranea* was used in all assays. Planarian colonies were maintained in clear, food-grade plastic containers holding 1,500–2,000 mL of 1× Montjuïch salted medium, kept in darkness at 20 °C. Planarians were not subjected to antibiotic treatment.

Colonies were fed fresh, strained beef liver purée once per week, cleaned twice weekly, and maintained at a population density of approximately 200–400 animals per container. For detailed colony preparation and long-term maintenance, see^32^.

##### Pre-experiment Conditioning

Prior to experimental infection, starve planarians for 7–12 days to standardize metabolic and size conditions. Use groups of 5–10 animals per well for each experimental condition. Ensure equal numbers of animals across all conditions, including positive and negative controls.

##### Animal Size Selection

- Select planarians ∼5 mm in length, corresponding to ∼ 5 x 10^5^ host cells per animal^54^.
- Planarians <5 mm in length can lead to inconsistent results, with planarians that have blastemas or incomplete development.
- Planarians >5 mm in length are less ideal for fluorescence immunostaining.
- Planarians ∼5 mm in length achieve reliable host outcomes within our planarian colony and were used for establishing our reported ID_50_ and LD_50_ values.

NOTE: To facilitate consistent size selection, print a representative 5 mm × 5 mm grid and place it inside a plastic binder sleeve beneath the transparent planarian container. Ensure planarians are fully extended before measuring length.

##### Health and Inclusion Criteria

**CRITICAL STEP:** Examine all planarians under a stereomicroscope prior to use.

**Exclude animals with visible damage or injury:**

- Presence of blastemas or incomplete pigmentation
- Abnormal pigmentation (significantly darker or lighter regions)
- Missing or damaged tissue `o` Visible lesions or scars from prior injury
- Open wounds or unhealed tissue `o` Abnormal body shape or deformity
- Any chronic or non-healing condition

**Exclude animals with developmental abnormalities:**

- Bifurcated or duplicated head/tail regions
- Misshapen or asymmetric body morphology
- Incorrect number of photoreceptors (only one bilateral pair is acceptable)
- Abnormally positioned pharynx
- Defective ciliation resulting in impaired motility or irregular gliding behavior

Inclusion criteria require animals to be of the correct length, fully developed, and free of injuries or physical or genetic abnormalities.

##### Plate Preparation

1. Transfer healthy planarians that meet all inclusion criteria into 2 mL of fresh planarian water per well in a non-tissue culture-treated 6-well polystyrene plate.
2. Assign wells for:

- Negative controls: Uninfected/mock animals.
- Positive controls: Animals infected with wild-type *C. albicans*.
3. Remove and replace all water to ensure uniformity across wells.

NOTE: Transfer animals to 6-well plate one day prior to infection to allow acclimation to the new environment and population density.

#### 3. Infection Setup (Figure 1)

##### Prepare Inoculation

1. Using aseptic technique, gently remove planarian water from each well of the 6-well plate.
2. Add the calculated volume of *C. albicans* culture to each well.
3. Bring the final volume to 2 mL per well with fresh planarian water.
4. Process one well at a time to ensure that planarians remain fully submerged at all times. NOTE: Avoid exposing animals to air, as transient drying or agitation can cause stress and alter infection outcomes.

##### Experimental Plate Incubation Conditions

1. Place the inoculated 6-well plate in a dark, static, and temperature-stable location (∼20-25 °C).
2. Record the date and time of inoculation (designated as time 0 h).
3. Maintain static incubation – do not shake, agitate or disturb plates.

NOTE: Avoid high-traffic or vibration-prone areas (e.g., drawers, instrument surfaces, or busy benchtops), as movement can stress the animals and confound results.

**Figure 1:**
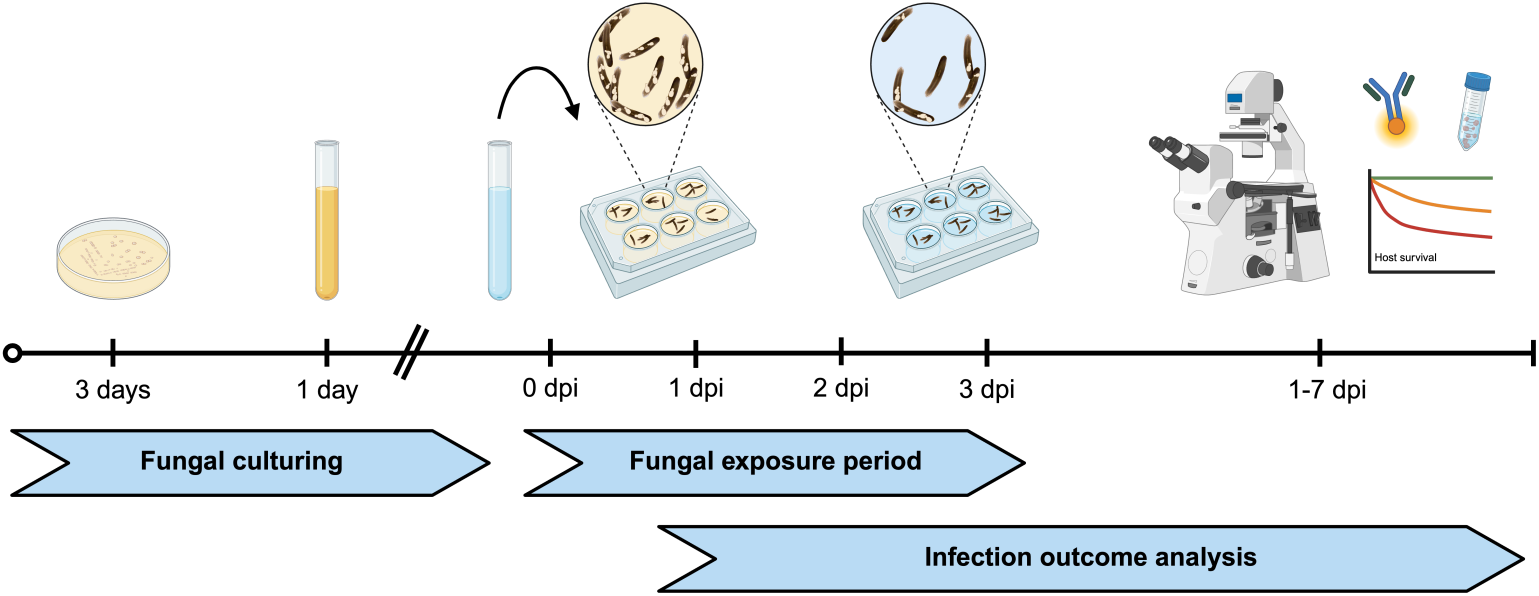
Experimental design for planarian fungal infection. *C. albicans* strains are cultured on YPD agar for three days, followed by a 16-h incubation in liquid YPD with shaking. Cells are harvested, washed in planarian water, quantified spectrophotometrically, and added directly to wells containing 5-10 planarians in planarian water for continuous exposure of up to 72 h. Host-pathogen outcomes are recorded every 24 h from 17 days post-infection (dpi). After the exposure period, animals are gently washed daily. Figure created with BioRender.

##### Resuspension of Fungal Cells

- At each timepoint, gently resuspend settled *C. albicans* cells without disturbing the planarians.
- At 24 h post-infection (hpi):

1. Using a sterile 3 mL transfer pipette, tilt the plate slightly toward you.
2. Aspirate the medium and gently dispense it back into an area of the well free of animals to mix the settled fungal layer.
3. Repeat this step 5-10 times per well, avoiding direct contact with the planarians.
4. Observe fungal settlement against a dark background (e.g., black benchtop or dark paper) for best visibility.
5. Resuspend uninfected control wells under identical conditions to maintain experimental consistency between groups.
- At 48 hpi:

6. Repeat the resuspension procedure described above.
7. Perform all resuspension steps within +/− 1 h of the designated timepoint to ensure consistency across replicates.

##### End of Exposure (72 hpi)

8. After 72 hpi, gently transfer planarians to a new 6-well plate containing 2 mL of fresh planarian water per well.
9. Minimize the transfer of fungal culture by using the least amount of medium when transferring the animals.
10. Remove and replace the old liquid with fresh planarian water.

NOTE: Handle animals and any tissue fragments gently to avoid mechanical injury or any additional stress.

#### 4. Host Endpoint Assessment

##### Recording Host Outcomes

Begin assessing host outcomes at 1-day post-infection (dpi) and continue daily until desired endpoint. Quantify host damage using a standardized 0–3 scoring system (**Figure 2**):

- 0: No change from uninfected controls (healthy, asymptomatic)
- 1: Mild symptom(s)
- 2: Severe symptom(s)
- 3: Death

**Figure 2:**
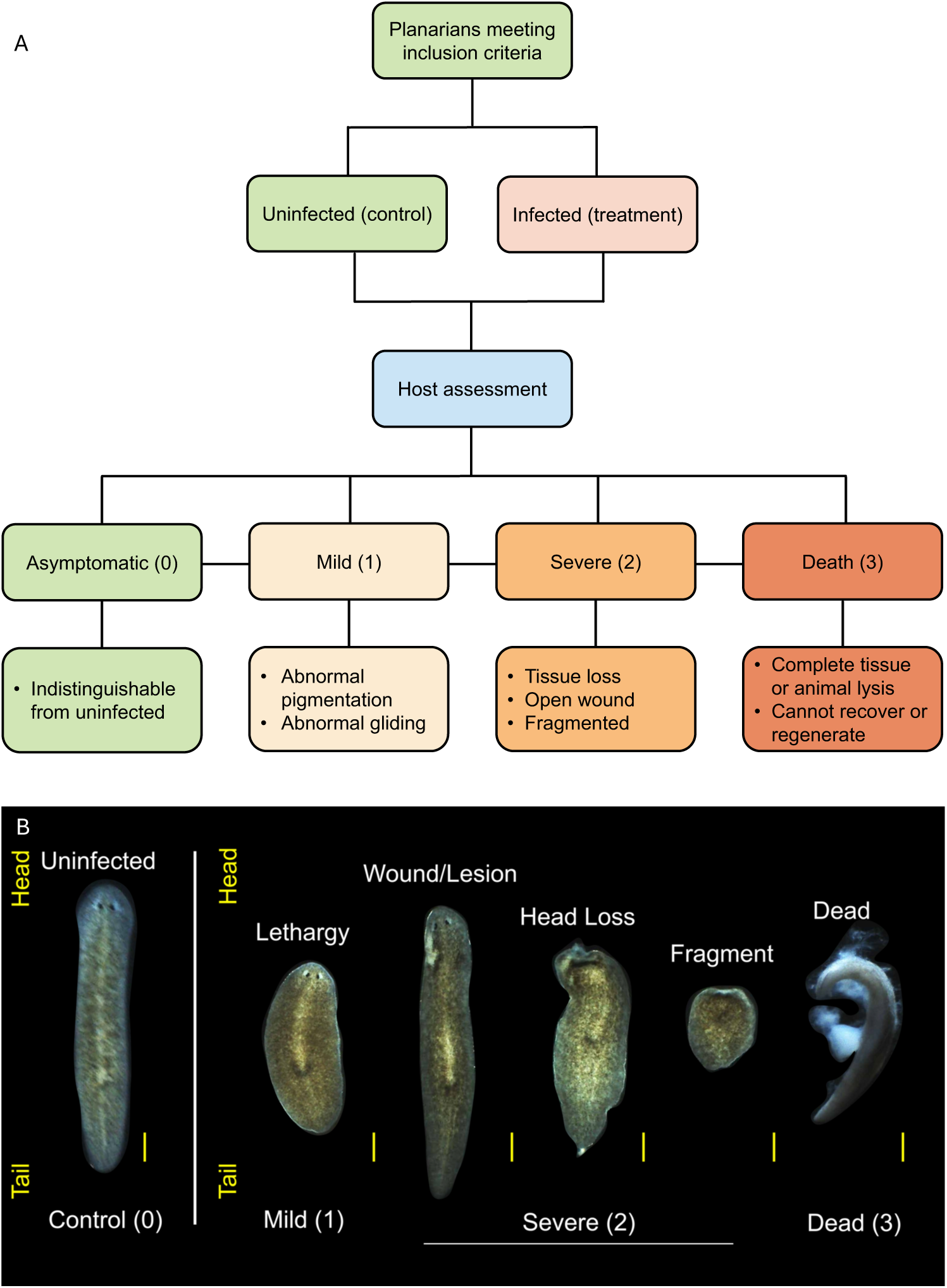
Standardized scoring of planarian host health during *C. albicans* infection. (A) Flowchart outlining the systematic scoring of planarian host outcomes on a 0-3 scale. (B) Representative host phenotypes at 4 dpi in uninfected planarians or planarians infected with 7.0×10^7^ cells/mL of wildtype *C. albicans* illustrating each score in the standardized health index: 0 = asymptomatic; 1 = mild symptoms; 2 = severe symptoms; 3 = host death. Scale bar, 200 µm.

##### Criteria for Host Symptom Evaluation

- **Score 0 (Asymptomatic):** Animals are either uninfected or infected with no observable changes; they display normal morphology, pigmentation, and responsiveness.
- **Score 1 (Mild):** At least one of the following is observed relative to asymptomatic controls:
  - Reduced speed or abnormal gliding
  - Partial or complete loss of phototactic response
  - Sustained or repetitive body contractions
  - Abnormal/non-uniform pigmentation (lighter or darker patches)
  - Small lesions or minor tissue regression (head/tail regression)
- **Score 2 (Severe)**: One or more of the following symptoms are present (animals remain alive):
  - Eye film or loss of one/both eyes
  - Complete head loss or severe anterior damage
  - Major tissue loss in the pre-pharyngeal region
  - Fragmentation into multiple discrete body regions (cephalic, pharyngeal, posterior)
  - Open wounds with or without mucus secretion or internal leakage
  - Partial tissue lysis (loss of structure or integrity)
  - “C-shape” body posture or partial or complete paralysis
- **Score 3 (Death)**: Defined by complete tissue or whole-animal lysis, total loss of response to stimuli, and/or failure to recover or regenerate.

NOTE: **Figure 2** provides representative images and features for each score to ensure consistent and systematic host evaluation.

##### Determining Host Survival After Fungal Exposure

1. At 3 dpi, gently transfer planarians to a new 6-well plate containing ∼2 mL of fresh planarian water. Avoid transferring residual fungal culture.
2. Observe animals using light microscopy (≥10× objective). Record all data using the standardized scoring sheet **(Supplemental File 2)**.
3. Determine live animals using one or more of the following criteria:
  - Negative phototaxis or response to gentle transfer pipette perturbation
  - Intact body structure
  - Stable attachment to the plate
4. Determine dead animals using one or more of the following:
  - Complete lysis or disintegration (residual debris only)
  - Transparent, eviscerated, or non-regenerating body
  - Absence of movement or recovery
5. Promptly remove dead animals and debris after viability confirmation under microscopy.

**KEY STEP:** Remove dead animals immediately to prevent cross-contamination or altered survival metrics. If debris or fungal carryover remains, transfer surviving animals again to a fresh plate with clean planarian water to prevent *C. albicans* reseeding or biofilm formation. NOTE: Place the 6-well plates over a dark background to enhance contrast during visual inspection steps.

#### 5. Fungal Burden Assessment

##### Colony Forming Unit (CFU) Quantification

1. At desired infection timepoints, collect planarians (at least 5 animals per condition per day) and include matched uninfected/mock controls.
2. Gently rinse animals with fresh planarian water two times to remove background C. albicans.
3. Transfer the planarians into labeled 1.7 mL or 5 mL Eppendorf tubes and remove water.
4. Add 300 μL planarian water per tube.
5. Homogenize animals using a sterile pestle until no large fragments remain (∼30-45 s).
6. Perform serial dilutions of the homogenate (1:10, 1:100, etc.).
7. Plate 100 μL of each dilution onto YPD agar plates supplemented with 50 μg/mL each of ampicillin, rifampicin, streptomycin, and neomycin. Spread using sterile glass beads or an inoculation stick.
8. Leave the plates on the bench for 5–10 min after spreading the homogenate to permit absorption into the agar and prevent liquid from dripping off the plate.
9. Prepare triplicate plates per dilution and condition (e.g., 3 plates for 1 dpi infected and uninfected animals).
10. Incubate plates at 30°C for 48 h.
11. Count CFUs, multiply by the dilution factor, and normalize to the number of planarians to calculate the average CFUs per planarian and condition.

#### 6. Immunofluorescence Anti-*Candida* Staining

NOTE: See **Supplemental File 1** for antibody details and reagent sources. Perform all steps with gentle orbital rocking (100-140 rpm).

1. Collect animals from each condition/timepoint and transfer into labeled 20 mL scintillation vials containing ∼2mL planarian water.
2. Gently remove all medium, rinse once with ∼2mL of fresh planarian water, remove water and replace medium.
3. Sacrifice animals in 7.5% N-Acetylcysteine (NAC) in 1× PBS for 3 min.
4. Gently remove liquid and fix animals in 4% formaldehyde in 0.3% PBSTx (PBS + Triton X-100 for 15 min.
5. Rinse twice with 1× PBS.
6. Permeabilize with 1% SDS for 15 min, then rinse three times with 1× PBS.
7. Bleach in 6% H₂O₂ in 1× PBS under a bright LED light for 4–12 h). NOTE: If bleaching cannot proceed immediately, store fixed samples in 1× PBS at 4 °C (for up to 1 week or dehydrate in 100% methanol for longer term storage).
8. Transfer animals to 24-well plates containing 1× PBS.
9. Block nonspecific binding in PBS-TB for 4 h at room temperature (or 8 h at 4°C)
10. Incubate with primary anti-*Candida* antibody (PA1-27158, Thermo Fisher; 1:500) for 4 h at room temperature (or 8 h at 4°C).
11. Remove primary antibody and wash 8 times every 20 min (2.5 h total) with 0.3% PBSTx.
12. Incubate with Alexa-568-conjugated anti-rabbit secondary antibody (A-11011, Thermo Fisher; 1:800) for 4 h at room temperature (or 8 hours at 4°C). NOTE: Keep samples covered to prevent photobleaching; antibodies can be reused.
13. Remove secondary antibody and wash every 8 times (2.5 h total) with 0.3% PBSTx.
14. Image using yellow-green excitation (∼578 nm) and observe orange-red emission (∼603 nm).

## REPRESENTATIVE RESULTS

We describe an updated approach for infecting planarians with *C. albicans* (**Figure 1**) and for systematically quantifying host outcomes (**Figure 2**). This refined workflow supports robust assessment of multiple infection parameters, including morbidity (symptomatic disease severity), survival (lethality), fungal burden (CFUs), and spatial-temporal dynamics of pathogen colonization via fluorescence immunostaining. Unlike most invertebrate infection models that primarily enable binary live/dead readouts, this planarian model captures a richer spectrum of infection outcomes and host responses.

Following fungal infection, hosts are classified into one of four outcome categories – asymptomatic, mild disease, severe disease or death – based on standardized criteria (**Figure 2A**). Updated representative phenotypes within each category, including distinct behavioral aberrations after infection (**Figure 2B**), provides reproducible and easily recognizable benchmarks for disease progression. This classification framework balances facilitates broad phenotypic capture while maintaining consistent and quantifiable endpoints.

Complementary assays – global fungal burden, morbidity/mortality scoring, and spatialtemporal imaging – collectively validate infection status and disease severity in a dosedependent manner. As shown in **Figure 3**, fungal colonization is markedly reduced, and by 7 dpi, nearly all surface-associated fungi are cleared. These observations demonstrate the rapid and effective innate immune response mounted by *S. mediterranea* as previously reported^23^.

**Figure 3:**
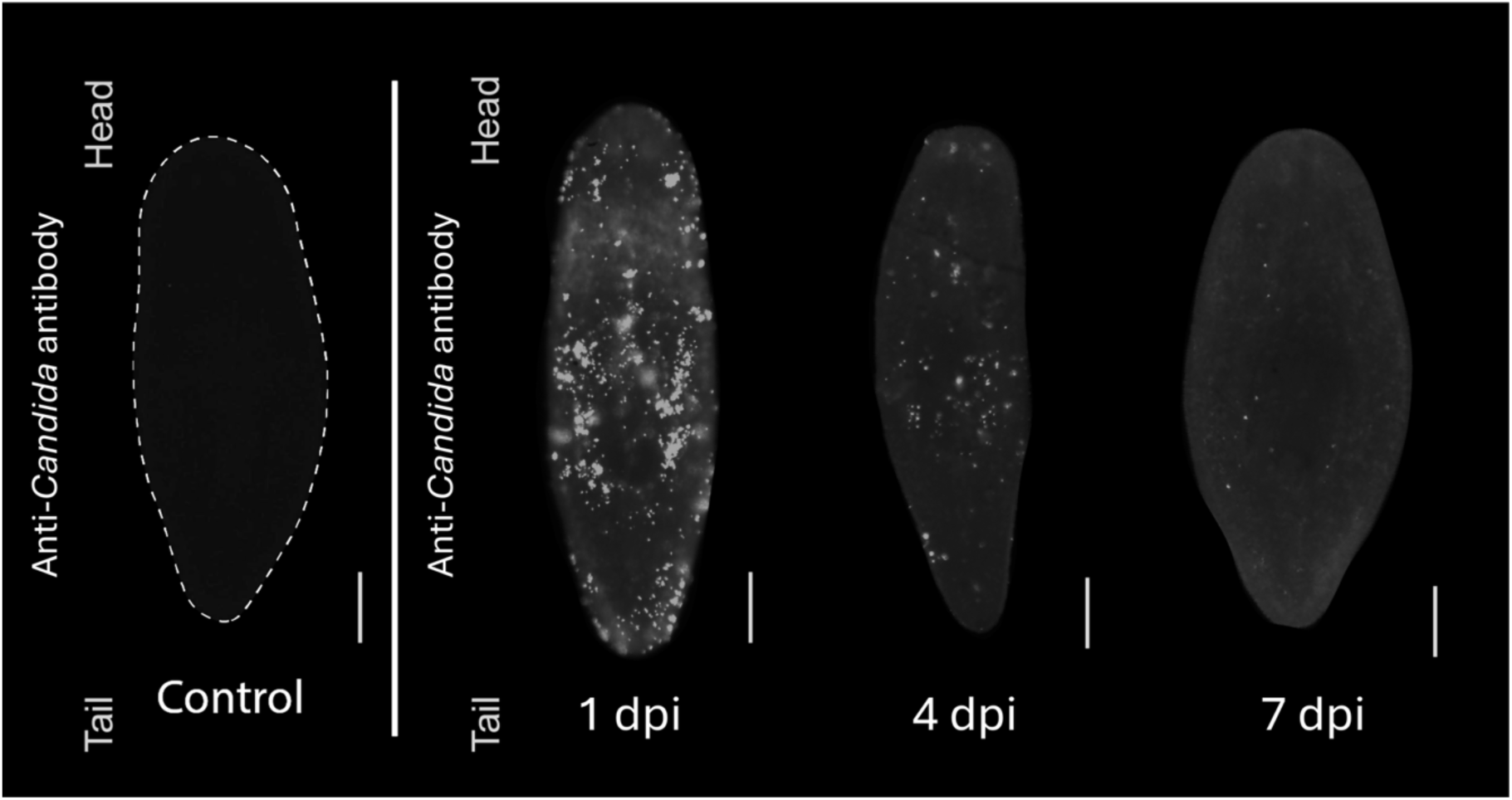
Immunofluorescent detection of *C. albicans* colonization on planarian epithelial surfaces. Planarians were infected with 7.0×10^7^ cells/mL of wildtype *C. albicans* and collected at 1, 4, and 7 dpi. Fungal cells were detected using an anti-*Candida* primary antibody and Alexa Fluor 568-conjugated secondary antibody. Fluorescence labeling denotes attached *C. albicans* cells on planarian epithelial surfaces. Scale bar, 200 µm.

In **Figure 4**, post-infection outcomes are quantified across exposure doses. Uninfected controls remain entirely asymptomatic. At a dose of approximately 7.0×10^7^ *C. albicans* cells/mL, approximately 30% of infected animals show no symptoms, 50% present with one or more symptoms, and 20% are dead. At a dose of approximately 8.0×10^7^ *C. albicans* cells/mL, 30% are asymptomatic, 20% are symptomatic, and 50% are dead. The difference in host outcomes between doses is statistically significant (isometric log-ratio (ILR)transformed MANOVA, * *p* = 0.04.

**Figure 4:**
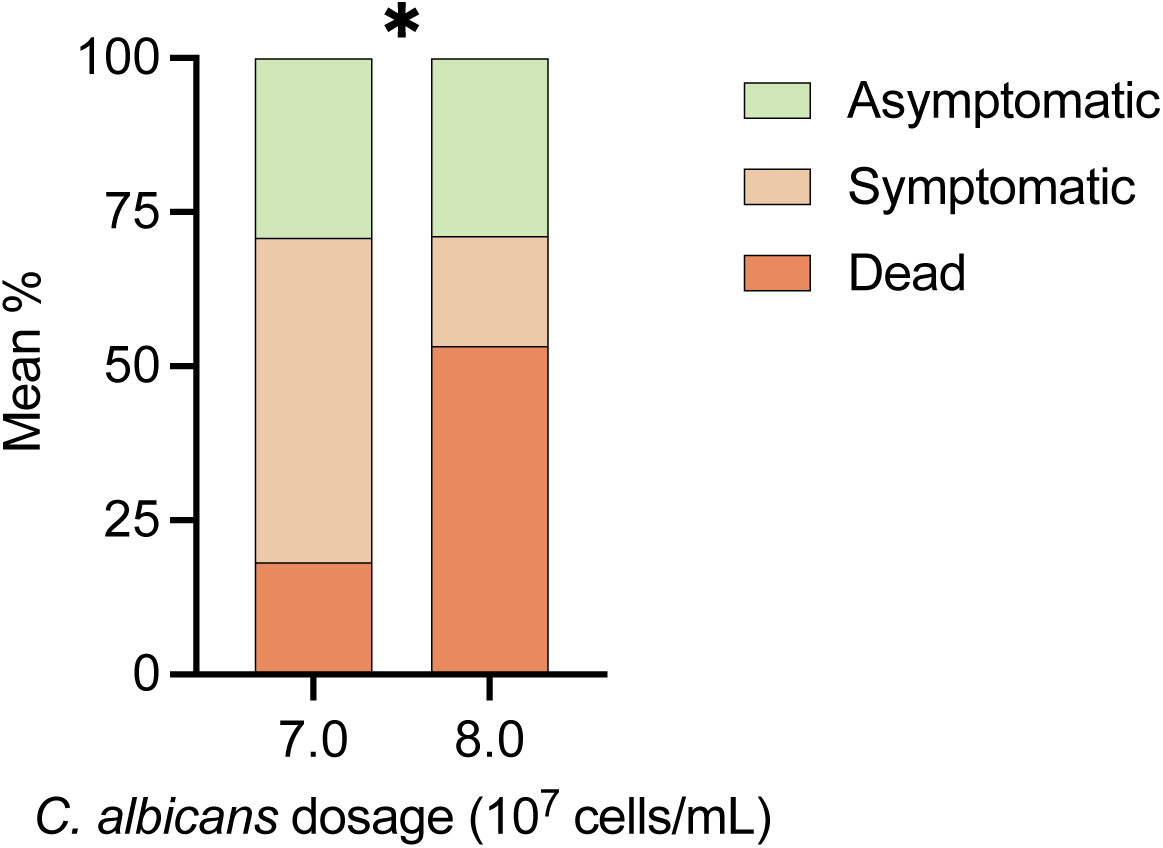
Categorical host outcomes following infection with wildtype *C. albicans*. Host outcomes were assessed at 4 dpi following exposure to 7.0×10^7^ cells/mL or 8.0×10^7^ cells/mL of wildtype *C. albicans*. Mockinfected controls exhibited no symptoms or deaths. Infected animals were categorized into 3 groups: asymptomatic (infected but phenotypically indistinguishable from uninfected animals), symptomatic (exhibiting one or more pathological symptoms) or dead. At least nine biological replicates (ten animals per replicate) were analyzed. Statistical differences in overall host health between infection doses were determined using isometric log-ratio (ILR) transformation with MANOVA; * *p* = 0.04.

**Figure 5** illustrates dose-dependent survival across increasing *C. albicans* concentrations (7.0-8.5×10^7^ cells/mL). Median survival declines from 90% at the 7.0×10^7^ cells/mL to 50% at 8.0×10^7^ cells/mL, and <25% at 8.5×10^7^ cells/mL. Kaplan-Meier survival analysis (**Figure 6**) further confirms an overall survival of 45% for planarians infected with the wildtype *C. albicans* approximate LD_50_ dose (8.0×10^7^ cells/mL), compared to an overall 100% survival in uninfected mock controls (95% confidence interval, 30-70%).

**Figure 5:**
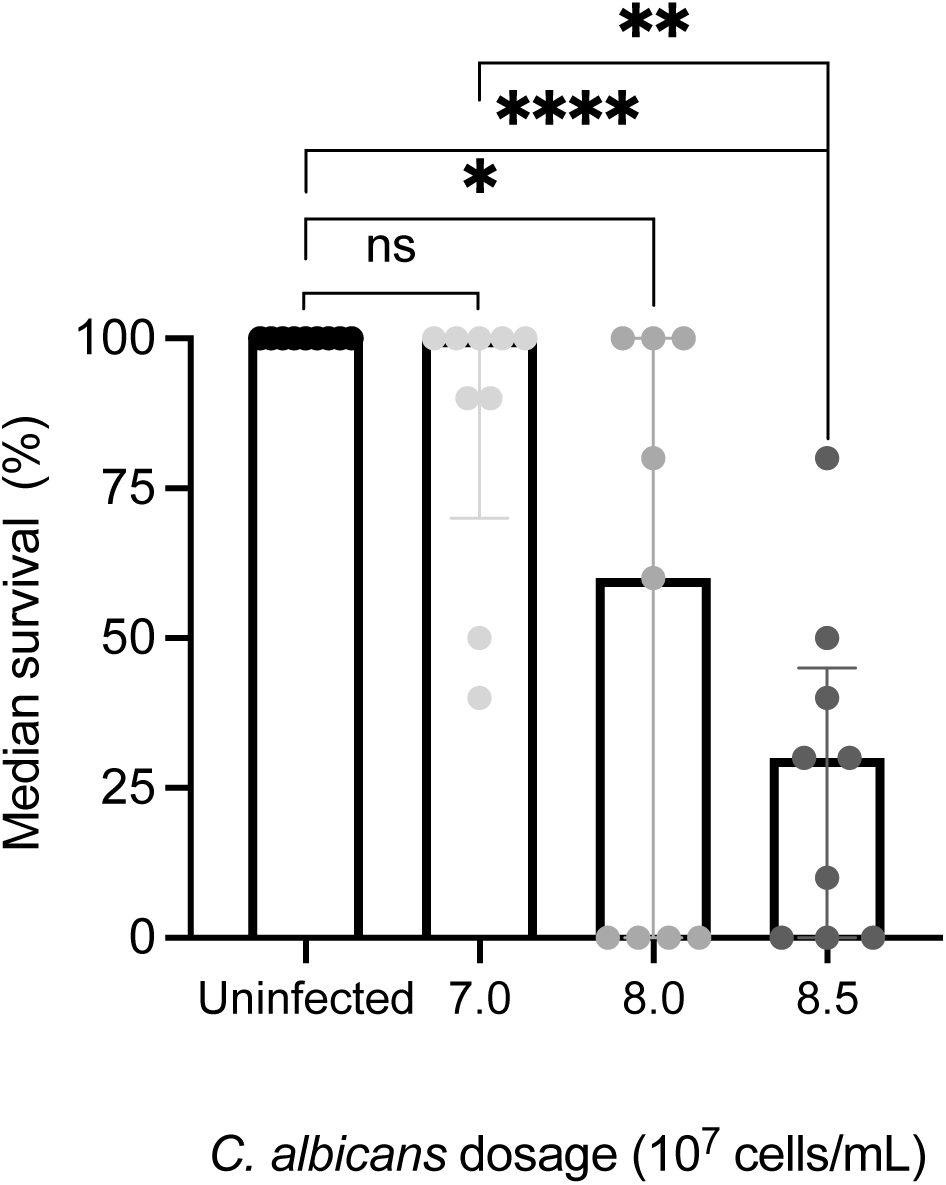
Host median survival decreases in a dose-dependent manner following infection with wildtype *C. albicans*. Median host survival was assessed across infection doses: 7.0×10⁷ cells/mL, 8.0×10⁷ cells/mL, and 8.5×10⁷ cells/mL of wildtype *C. albicans*. Uninfected controls exhibited 100% survival. Each data point represents five planarians (for the 8.0×10⁷ cells/mL dose) or ten planarians (for the 7.0×10⁷ cells/mL and 8.5×10⁷ cells/mL doses). Nine replicates were performed across at least 45 animals per group. The interquartile range is represented for each condition. Asterisks indicate levels of statistical significance; ns = no significance; * *p* < 0.05; ** *p* < 0.01; *** *p* < 0.001; **** *p* < 0.0001, determined using single-tailed Kruskal-Wallis and Dunn’s post hoc tests with Bonferroni correction.

**Figure 6:**
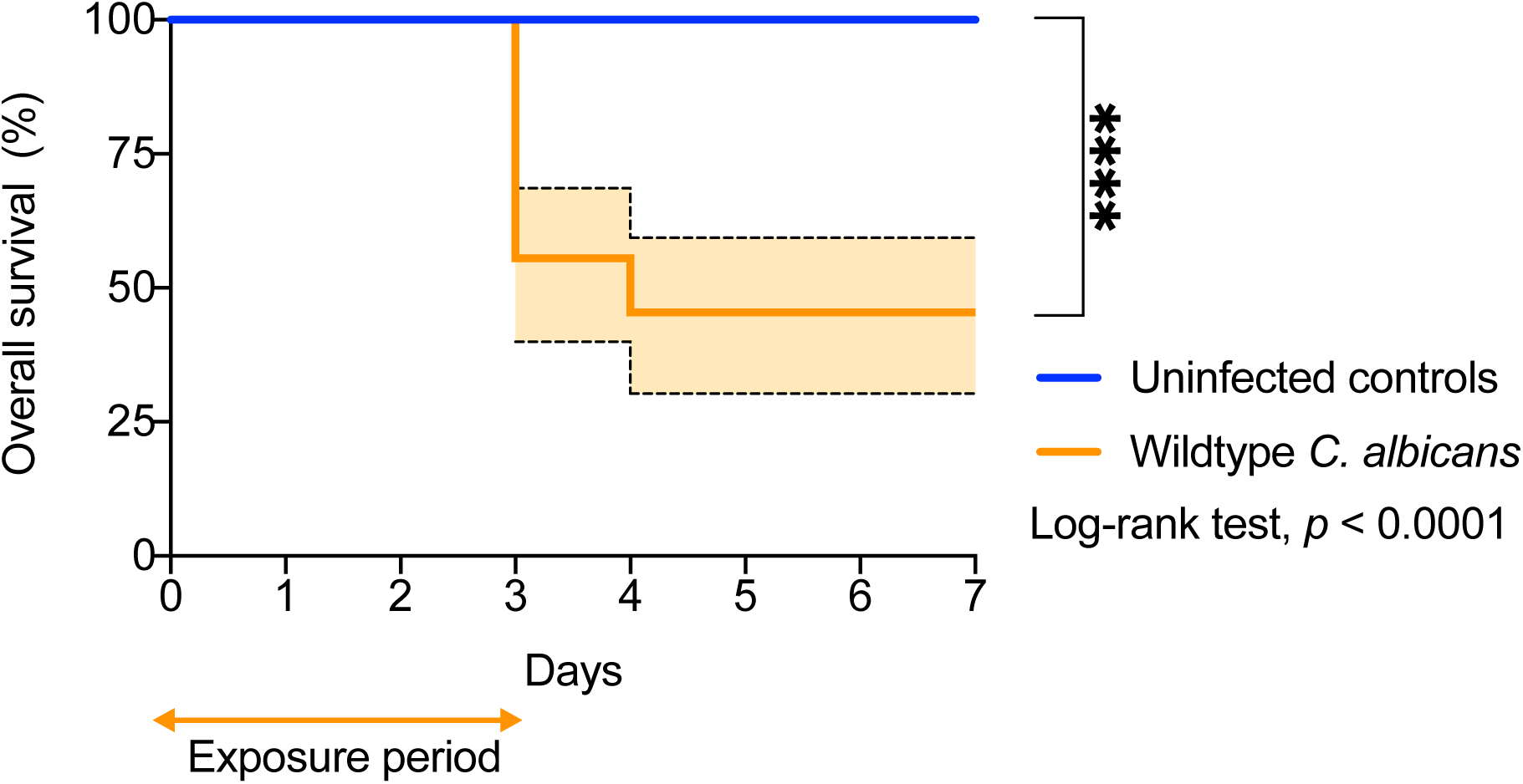
Kaplan-Meier survival analysis of planarians infected with wildtype *C. albicans*. Kaplan-Meier survival curves compare mock infected planarians with those infected with wildtype *C. albicans* (8.0×10⁷ cells/mL; LD_50_). Overall survival was 100% for uninfected controls and 45.5% for infected animals (95% confidence interval, 30 to 70%). Survival was significantly lower in infected animals compared to controls, determined using Mantel–Cox log-rank test. **** *p* < 0.0001. A total of 45 animals per group were analyzed across nine replicates, of which five were biological replicates (5 animals per replicate). All planarians were maintained under identical conditions and monitored for 7 days total following a 3-day exposure period.

Together, these results establish reproducible, quantifiable endpoints for analyzing fungal pathogenesis and host defense in the *S. mediterranea* model.

## DISCUSSION

Here, we demonstrate that varying the infectious dose of wildtype *C. albicans* produces reproducible, dose-dependent differences in morbidity in *S. mediterranea*, underscoring the model’s sensitivity for quantitative assessment of fungal virulence. We outline strategies for establishing baseline infectious and lethal doses within laboratory-maintained planarian colonies and present criteria for confirming successful systemic infection via soaking, integrating both host health scoring and measurements of fungal burden using CFUs and immunostaining.

Through systematic optimization, we identified several experimental parameters that are essential for achieving consistent infection dynamics and reproducible host outcomes. Beyond inoculum concentration, factors such as planarian size, population density, and surface area-to-volume ratio markedly influence the progression and severity of infection. The refinements introduced here – including standardized inclusion criteria, controlled infection conditions, and detailed endpoint scoring – substantially improve reproducibility and enable meaningful comparisons across experiments and laboratories.

Planarians offer a rare combination of experimental accessibility and biological complexity, allowing *in vivo* investigation of host-pathogen interactions in a multicellular organism. Their size facilitates rapid macroscopic assessment of disease phenotypes, including motility defects, morphological changes, and survival, while whole-mount immunostaining provides cellular-resolution visualization of host responses throughout the body. Unlike cell culture systems, the planarian model allows real-time analysis of fungal engagement with multiple tissues in an intact organism. Moreover, this protocol supports multisystemic analyses, enabling the study of coordinated responses across epithelial, neural, and immune compartments. These advantages – combined with low cost, minimal ethical constraints, and compatibility with high-throughput screens – establish planarians as a powerful and practical alternative host for studying fungal pathogenesis.

The planarian model also has inherent limitations. The absence of an adaptive immune system precludes analysis of T-cell- and B-cell-mediated processes and prevents investigation of adaptive immunological memory. Physiological differences from mammalian hosts – including body temperature, tissue architecture, and organ organization – limit the degree to which certain aspects of mammalian disease progression can be recapitulated. Additionally, planarians’ robust regenerative capacity, while highly valuable for studying tissue repair, can complicate interpretation of infection phenotypes, as accelerated wound healing may obscure the full extent of pathogen-induced damage.

A major strength of the soaking infection protocol is that it enables simultaneous assessment of pathogen behavior and host responses beginning at the earliest stages of colonization and mimics the most common route of human fungal pathogenesis – epithelial overgrowth and barrier disruption. Soaking also facilitates simple and rapid onset and termination of the fungal challenge period, as media containing *C. albicans* can be simply added and removed from the wells. On the pathogen side, the system supports quantitative evaluation of colonization, tissue penetration, and lethality across diverse doses, strains, and conditions. On the host side, planarians permit accessible biochemical, cellular, and genetic analyses to examine epithelial barrier integrity, wound healing, stress responses, and innate immune activation. The platform is also readily scalable, cost-effective, and fully compatible with RNAi-mediated gene perturbation, enabling systematic interrogation of host pathways involved in defense, regeneration, and tissue homeostasis^35,36,55^.

Overall, this protocol enables a broad range of applications, including high-throughput screening of pathogen virulence, testing of mutant strain libraries, evaluation of antimicrobial activity, comparative studies across microbial species, and comprehensive assessment of morbidity and mortality in a live host. It also supports detailed analysis of conserved innate immune pathways and regenerative mechanisms. Time-resolved monitoring of infection and recovery allows fine-scale examination of how tissues resist, respond to, and repair following fungal challenge. Compared with other animal models of infectious disease, planarians occupy a complementary and valuable niche – ethically favorable, experimentally versatile, and biologically informative. As such, this platform serves as an effective bridge between reductionistic *in vitro* systems and more complex vertebrate models, facilitating discovery of both fungal pathogenic strategies and host protective mechanisms.

## Supporting information

Supplemental File 1 (Media and Buffers Recipes)

Supplemental File 2 (Host Outcomes Tracking Template)

## ACKNOWLEDGMENTS

We thank all members of the Nobile, Oviedo, and Hernday labs for insightful discussions about the *C. albicans*–*S. mediterranea* infection model. We are grateful to Edelweiss Pfister for lab management and planarian maintenance. This work was supported by the National Institutes of Health (NIH) National Institute of General Medical Sciences (NIGMS) awards R35GM156045 and R35GM124594 to C.J.N., and R35GM158501 and R01GM132753 to N.J.O. This work was also supported by the Kamangar family in the form of an endowed chair to C.J.N. N.M.S. was supported by the Center for Cellular and Biomolecular Machines (CCBM) National Science Foundation (NSF) Center of Research Excellence in Science and Technology (CREST) fellowships under awards NSF-HRD-1547848 and NSF-EES-2112675. The content is the sole responsibility of the authors and does not represent the views of the funders. The funders had no role in the design of the study, in the collection, analysis, or interpretation of data, in the writing of the manuscript, or in the decision to publish the results. N.M.S. acknowledges the use of ChatGPT for assistance with editing the manuscript (checking spelling, grammar, formatting, and sentence structure).

