## Supplemental File 1 (Media and Buffers Recipes) for "An optimized protocol for *Candida albicans* infection in *Schmidtea mediterranea* to study fungal pathogenesis and host defense"

### **YPD *C. albicans* growth medium**

1% yeast extract, 2% peptone, 2% glucose; pH 6.8

1. Add 10 g of yeast extract
2. Add 20 g of peptone
3. Add 20 g of dextrose
4. Add 20 g of agar (for plates only)
5. Dissolve the reagents in distilled, deionized water and adjust the final volume to 1 L
6. Autoclave for 20 min at 121°C and 15 psi using “liquid” cycle

**Planarian water/medium (1x Montjuïch salts)**

**Sodium chloride (NaCl) – 1.6 mM, Calcium chloride dihydrate (CaCl₂·2H₂O) – 1.0 mM, Magnesium sulfate heptahydrate (MgSO₄·7H₂O) – 1.0 mM, Magnesium chloride hexahydrate (MgCl₂·6H₂O) – 0.1 mM, Potassium chloride (KCl) – 0.1 mM, Sodium bicarbonate (NaHCO₃) – ~1.2 mM; pH 7.5**

1. **Add 93 mg Sodium chloride (NaCl)**
2. **Add 147 mg Calcium chloride dihydrate (CaCl₂·2H₂O)**
3. **Add 246 mg Magnesium sulfate heptahydrate (MgSO₄·7H₂O)**
4. **Add 20 mg Magnesium chloride hexahydrate (MgCl₂·6H₂O)**
5. **Add 7.5 mg Potassium chloride (KCl)**
6. **Add 101 mg Sodium bicarbonate (NaHCO₃)**
7. Dissolve the reagents in distilled, deionized water and adjust the final volume to 1 L
8. Filter sterilize by passing solution through a 0.22 μm filter
9. Media should be prepared weekly as pH drifts and salts precipitate out of solution
10. Store at room temperature

**Broad-spectrum antibiotic medium**

**Neomycin, streptomycin, rifampicin and ampicillin (50 μg/mL each)**

1. **Diluted each antibiotic stock to be 50 mg/mL working concentration**
   a. **NOTE: Rifampicin should be dissolved in 1% DMSO and protected from light degradation. All other antibiotics can be prepared using standard protocols**
2. **Prepare YPD-agar medium as previously described minus volume of antibiotic stocks**
3. **Following autoclaving, cool medium to 50-55**°C then add antibiotic stocks
4. Pour plates and store in 4°C in darkness. Plates are stable for 2 weeks

**Immunostaining reagents**

**10x Phosphate-Buffered Saline (PBS)**

1. Add 80 g sodium chloride (NaCl)
2. Add 2 g potassium chloride (KCl)
3. Add 14.4 g sodium phosphate dibasic (Na₂HPO₄)
4. Add 2.4 g potassium phosphate monobasic (KH₂PO₄)
5. Adjust pH to 7.4 and autoclave

**PBS with Triton X-100 (0.3% PBSTx)**

1. Add 3 mL Triton X-100
2. Dissolve in 997 mL of 1x PBS

**7.5% N-Acetyl-L-cysteine (NAC)**

1. Add 0.75 g NAC powder (store at 4°C)
2. Dissolve in 9 mL MilliQ water
3. Add 1 mL of 10x PBS

**4% Formaldehyde (FA)**

1. Add 1.1 mL of 35.6% formaldehyde
2. Dissolve in 8.9 mL of 0.3% PBSTx

**1% Sodium Dodecyl Sulfate (SDS)**

1. Add 1 mL of 10% SDS
2. Dissolve in 9 mL of 1x PBS

**6% Hydrogen Peroxide (H₂O₂)**

1. Add 2 mL of 30% H₂O₂ (store at 4°C)
2. Dissolve in 8 mL of 1x PBS

**PBS + Triton X-100 (PBSTx)**

1. Add 0.3 mL Triton X-100 per 100 mL of 1x PBS

**2.5% PBSTx with BSA (for blocking)**

2. Add 250 mg Bovine Serum Albumin (BSA)
3. Dissolve in 10 mL PBSTx
4. Filter sterilize by passing solution through a 0.22 μm filter
5. Store at 4°C

**1% PBSTx with BSA (for antibody dilutions)**

1. Add 100 mg Bovine Serum Albumin (BSA)
2. Dissolve in 10 mL PBSTx
3. Filter sterilize by passing solution through a 0.22 μm filter
4. Store at 4°C

**Anti-*Candida* primary antibody (PA1-27158, Thermo Fisher)**

1. Add 10 µL Anti-*Candida* antibody from stock tube
2. Dissolve in 5 mL filtered 1% BSA (working dilution is 1:500)
3. Store at 4°C

**Alexa Fluor 568 secondary antibody (A-11011, Thermo Fisher)**

1. Add 5 µL Alexa Fluor 568
2. Dissolve in 4 mL filtered 1% BSA (working dilution is 1:800)
3. Store at 4°C covered in foil
